# GFMBench-API: A Standardized Interface for Benchmarking Genomic Foundation Models

**DOI:** 10.64898/2026.02.19.706811

**Authors:** Ariel Larey, Elay Dahan, Amit Bleiweiss, Raizy Kellerman, Guy Leib, Omri Nayshool, Dan Ofer, Tal Zinger, Dan Dominissini, Gideon Rechavi, Nicole Bussola, Simon Lee, Shane O’Connell, Dung Hoang, Marissa Wirth, Alexander W. Charney, Yoli Shavit, Nati Daniel

## Abstract

The rapid scaling of Genomic Foundation Models (GFMs) has created a critical need for standardized evaluation frameworks. Current benchmarking practices are often fragmented, relying on model-specific preprocessing and inconsistent metric implementations that hinder reproducible comparisons. We present GFMBench-API, a high-level Python interface designed to unify the evaluation lifecycle of GFMs. GFMBench-API provides a modular “middleware” architecture that decouples model-specific tokenization and embedding logic from task-specific data streams and performance metrics. By standardizing the input/output schemas for common genomic tasks, such as regulatory element prediction, variant effect scoring, and long-range interaction mapping, GFMBench-API enables researchers to integrate new models or tasks with minimal “glue code.” Our interface ensures mathematical consistency across evaluations, providing a robust foundation for the transparent and systematic benchmarking of GFMs.

## Introduction

The rapid emergence of Genomic Foundation Models (GFMs) is transforming our ability to model complex biological sequences (1, 2), yet the field lacks a cohesive infrastructure for systematic evaluation. Current benchmarking efforts are frequently fragmented, with researchers forced to develop custom-made, non-interoperable pipelines to bridge the gap between specific model architectures and diverse biological and clinical tasks. This fragmented landscape is characterized by a wide, non-standardized formulation of tasks across disparate benchmarks, such as Genome Understanding Evaluation (GUE) (3), BEND (4), Nucleotide Transformer (NT) (5) benchmarks or tasks derived from the ClinVar dataset (6), which prevents meaningful cross-study comparisons.

Furthermore, the lack of a unified interface complicates the evaluation of a diverse range of task types, including supervised classification and zero-shot variant effect prediction (VEP) utilizing reference and alternative sequence inputs. Variations in how models handle conditional dependencies or specific paradigms, such as auto-regressive compared to masked language modeling (MLM) often lead to biased results depending on the task’s specific formulation.This creates significant technical debt, where codebases are dominated by ad-hoc scripts (“glue code”) patching together incompatible data formats and reimplementing methods (e.g. embedding distances) rather than core analysis. Significant engineering effort is thus wasted on re-implementing data loaders, sequence encodings, and evaluation protocols for every new model iteration.

To address these challenges, we introduce GFMBench-API, a universal middleware for the genomic machine learning ecosystem (Figure 1). By providing a standardized interface to connect model backbones with diverse genomic tasks, it shifts the focus from manual pipeline integration to robust, transparent comparison. Unlike existing studies that couple evaluation logic to specific models, GFMBench-API resolves infrastructural fragmentation through a unified abstraction layer. A detailed comparison to existing benchmarking efforts is provided in Appendix, Supplementary Note 1.

**Fig. 1.**
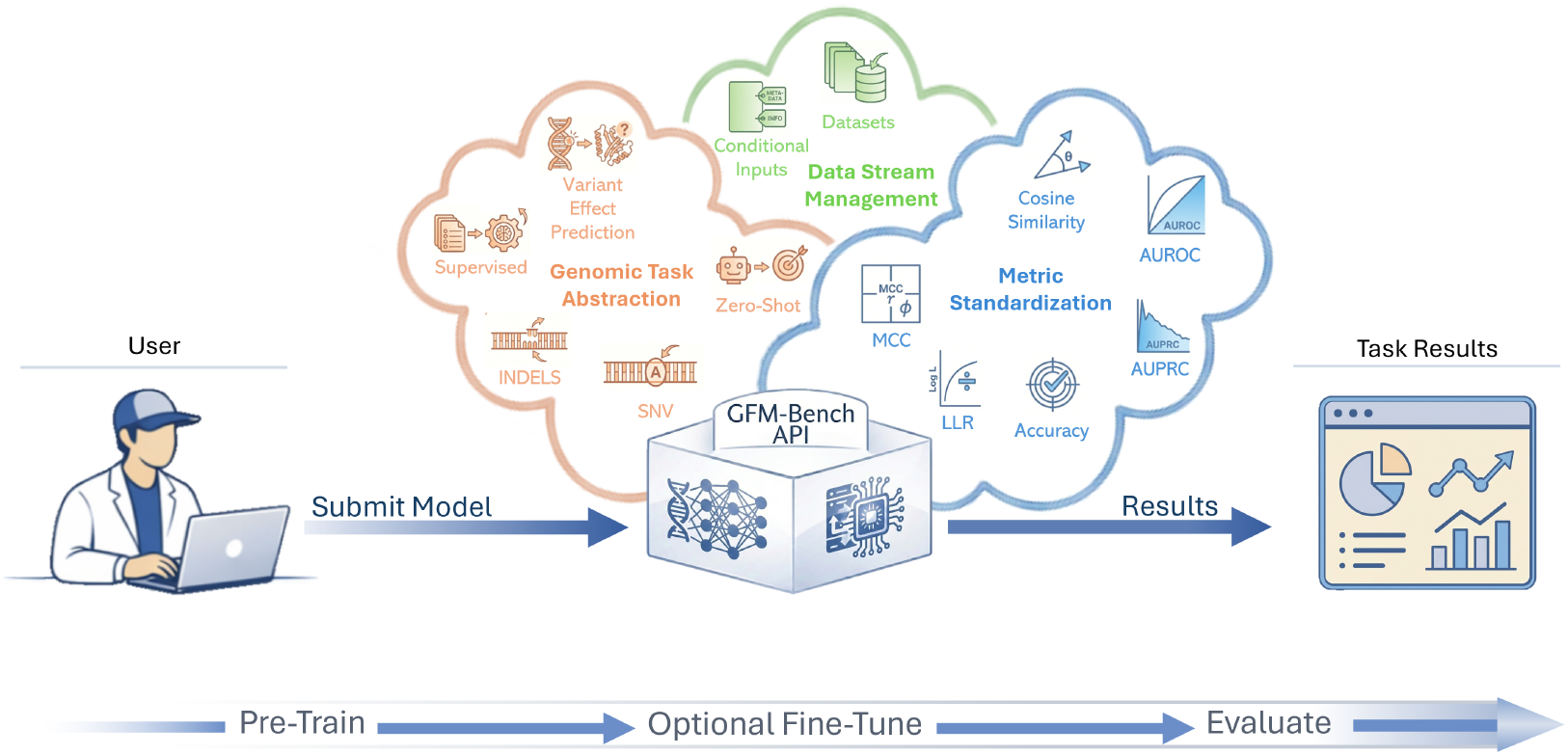
GFMBench-API offers a unified middleware that decouples genomic foundation model architectures from diverse tasks, standardizes data streams and metrics, and enables users to submit models through a single interface for reproducible evaluation and reporting.

In summary, our key contributions are as follows:

- We introduce an API middleware which decouples the model logic (tokenizer, inference etc.) from task execution, allowing any GFM to be evaluated across diverse benchmarks without custom “glue code”.
- Our proposed interface establishes a unified protocol for handling genomic inputs and outputs (including reference/alt allele handling and conditional logic), ensuring that disparate models are tested against identical genomic contexts and data streams.
- By centralizing metric implementation, we ensure that results are mathematically consistent, preventing metric drift and ensuring reproducibility.

## Technical Methods

GFMBench-API is a modular Python library that bridges diverse genomic architectures and standardized evaluation. Its extensible interface allows users to register tasks via a unified API, supporting scalable, distributed execution of models across a benchmark suite. Each task adheres to a shared interface containing its own datasets (train, validation, and holdout) and domain-specific metrics.

### A. Design Principles

The design of GFMBench-API is guided by a set of principles intended to ensure consistent, extensible, and model-agnostic evaluation of GFMs.

- **Separation of model development and task evaluation:** GFMBench-API separates model or algorithm development explicitly from task-level evaluation. Model development, including architectural choices, training procedures, and fine-tuning strategies, is performed by the user outside the scope of the task. This process uses information provided by the task such as task attributes and optional fine-tuning datasets. In contrast, task evaluation is fully encapsulated within the benchmark and executed using standardized datasets and metrics. This separation allows users full flexibility in how models are developed, while ensuring that evaluation is conducted in a strictly controlled and consistent manner, enabling fair comparisons across models.
- **Inference-driven metric compatibility:** Evaluation metrics in genomics are often applicable only to specific classes of models or training objectives. For example, masked likelihood–based metrics are appropriate for models supporting conditional probabilities over tokens under masking patterns (e.g., models trained with masked language modeling objectives), while causal sequence likelihood–based metrics are meaningful for autoregressive models that provide per-token probability estimates. To accommodate this variety, GFMBench-API defines a set of standardized inference method APIs at the model level. Developers may implement or wrap only the inference methods relevant to their algorithm, independently of any task implementation. During evaluation, tasks automatically compute only metrics compatible with the model’s inference outputs, ensuring both flexibility and correctness without requiring task-specific model adaptations.
- **Hierarchical task abstraction and reuse:** While all tasks conform to a shared base API, the variety of genomic evaluation settings naturally gives rise to groups of task types with common structure, such as supervised vs. unsupervised tasks, or classification vs. variant effect prediction tasks involving reference and alternate sequences. GFMBench-API captures this structure through hierarchical base task classes that define shared attributes and methods for each task category. Concrete tasks are implemented by extending these base classes and specifying only task and data-specific logic, reducing implementation complexity and promoting consistency across the benchmark.

### B. Implementation

The implementation of GFMBench-API reflects the design principles outlined above and provides a concrete realization of the proposed evaluation framework. We first present the Task API (subsection B.1), followed by a description of the hierarchical Task Design (sub-section B.2) and the Benchmark Tasks (subsections B.3) included in the current release.

#### B.1. Task API

The Task API provides a unified and model-agnostic interface through which users evaluate GFMs on extrinsic tasks, while preserving the separation between model development and task evaluation (Figure 2). Given a model to be evaluated, the user first initializes a specific task instance, after which all interactions with the benchmark proceed through a shared task API.

**Fig. 2.**
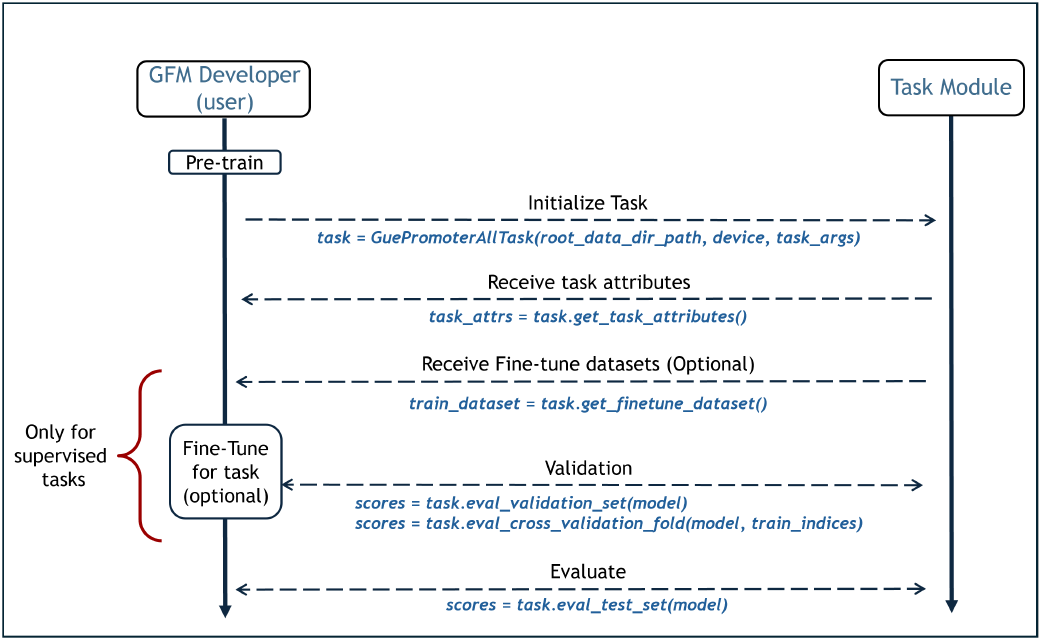
Overview of the Task API workflow for evaluating genomic foundation models. Users initialize tasks, optionally fine-tune models, and perform validation and test evaluation through a unified interface.

Upon initialization, users may query task metadata via the `get_task_attributes()` method, which returns a dictionary describing key properties of the task, such as whether it is supervised or zero-shot, the number of target classes, or whether the task involves variant effect prediction. These attributes inform downstream model development decisions without coupling the model implementation to the task itself.

For zero-shot tasks that do not require model adaptation, evaluation is performed directly by calling `eval_test_s et(model)`. This function returns performance scores for all evaluation metrics supported by the task and compatible with the model’s implemented inference methods. For supervised tasks, users may optionally obtain task-specific training data through `get_finetune_dataset()` and perform fine-tuning or task-specific adaptation externally. For instance, in a multi-class classification scenario, a user might map GFM sequence embeddings to label logits by plugging the embedding vector into a small trainable project layer. The resulting wrapped model is then passed to the task for evaluation.

During model development, users may evaluate intermediate performance by invoking `eval_validation_set(mo del)` when a validation split is available. If a predefined validation set is not available, the API enables user-defined cross-validation by allowing evaluation on separate folds via `eval_cross_validation_fold(model,fold_i ndices)`. In all cases, task execution, dataset handling, and metric computation are managed internally by the task implementation, ensuring standardized and reproducible evaluation across models and tasks.

Each task evaluates the model using multiple related metrics, with each metric requiring specific model outputs. To support this, users may implement any subset of a standardized collection of inference methods provided by the API, and only the methods implemented by the model are utilized during evaluation. The available inference methods support sequence-level, token-level, and variant-aware predictions, producing probabilities and/or embeddings as required (See Appendix, Supplementary Note 3 for more details). Tasks invoke only the methods needed for their metrics, with positional mappings handled automatically where tokenization affects alignment. This modular design enables flexible, consistent evaluation across diverse model architectures.

#### B.2. Tasks Design

The hierarchical task abstraction described above is implemented via a multi-level class hierarchy (Figure 3). At the root, `BaseGFMTask` defines the abstract task interface and shared infrastructure, including configuration parsing, dataset management, and the evaluation entry points described in the previous section. All tasks inherit from this class and implement its abstract methods: `get_task_name()`, `_create_datasets()`, `_get_default_max_seq_len()`, and `_eval_d ataset()`. Task attributes exposed through the API include indicators for the availability of fine-tuning and validation data, whether the task involves variant effect prediction, the number of target labels for classification, and an optional metadata schema describing conditional inputs required by the task (Appendix, Supplementary Note 4).

**Fig. 3.**
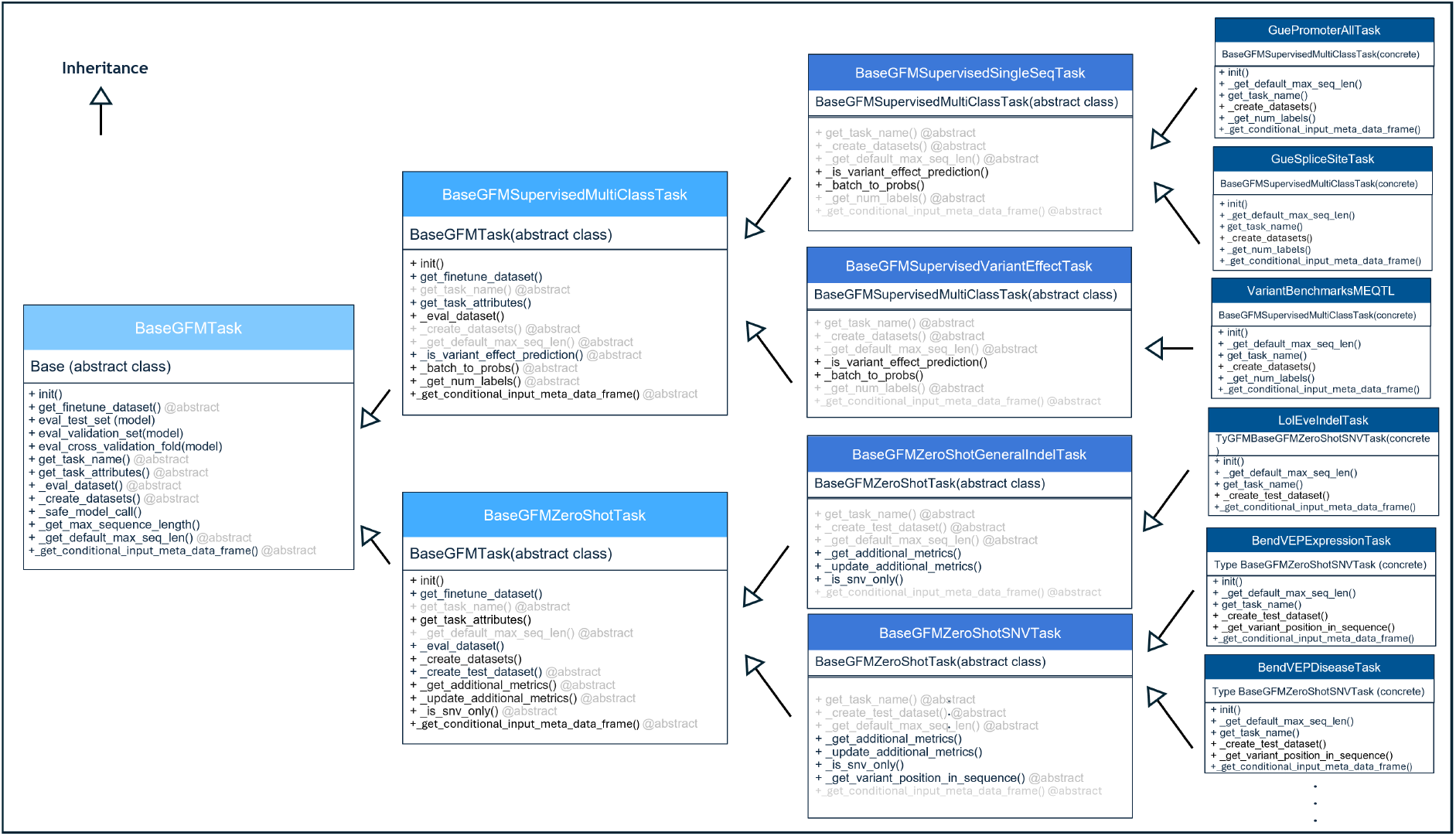
GFMBench-API class hierarchy. The hierarchy distinguishes supervised classification and zero-shot variant effect prediction tasks, with further specialization for different input representations. Concrete tasks inherit standardized evaluation behavior, ensuring consistent and reproducible benchmarking. Methods shown in black are implemented in the corresponding class, while methods shown in gray are inherited from or implemented in parent or child classes.

The hierarchy branches according to the evaluation paradigm. `BaseGFMSupervisedMultiClassTask` implements the evaluation loop for multi-class classification and computes a standard set of metrics, including classification accuracy, Matthews correlation coefficient (MCC), area under the receiver operating characteristic curve (AUROC), and area under the precision–recall curve (AUPRC). Two further specializations handle distinct data representations: `BaseGFMSupervisedSingleSeqTask` corresponds to the standard classification setting with a single DNA sequence as input, where models map sequences to class probability distributions; and `BaseGFMSupervisedVaria ntEffectTask`, which operates on paired reference and alternate sequences to enable supervised prediction of variant effects conditioned on both genomic contexts.

The zero-shot branch, rooted at `BaseGFMZeroShotTask`, implements evaluation using pre-trained model outputs without fine-tuning. All zero-shot tasks operate on paired reference and variant sequences and compute a common set of metrics, including log-likelihood ratio (LLR) derived from per-token sequence probabilities, as well as embedding-based similarity measures such as cosine similarity and L2 distance computed over sequence-level representative embeddings. Each metric is reported using both AUROC and AUPRC for binary pathogenicity classification. Two sub-classes further specialize this evaluation based on variant type: (a) `BaseGFMZeroShotSNVTask` incorporates position-specific metrics for single nucleotide variants, including cosine similarity computed at the variant-position embedding and masked token prediction LLR, which compares the model’s predicted probabilities for the reference and alternate alleles at a masked variant site; (b) `BaseGF MZeroShotGeneralIndelTask` supports arbitrary insertions and deletions using only sequence-level position-agnostic metrics, as localized comparisons are not well defined for length-altering variants.

Concrete tasks extend specific intermediate classes by implementing minimal methods: dataset construction, naming, sequence length, and label counts. They can optionally include metadata schemas for conditional inputs like tissue or cell types. See Appendix, Supplementary Note 2 for metrics’ details.

#### B.3 Benchmark Tasks

We demonstrate the usage of our API by implementing a diverse task benchmark, drawn from established genomic evaluation resources.

##### Supervised single-sequence tasks

Three tasks are derived from the GUE benchmark (3): promoter prediction, a binary classification task over 300 bp sequences; splice site prediction, a three-class classification task distinguishing donor, acceptor, and non-splice regions within 400 bp windows; and transcription factor binding site prediction, a binary classification task over 100 bp sequences aggregated from multiple ChIP-seq experiments. All GUE tasks provide predefined training, validation, and test splits.

##### Supervised variant effect tasks

Six tasks are derived from VariantBenchmarks (7): The coding pathogenicity task classifies coding-region single nucleotide variants (SNVs) as benign or pathogenic, the non-coding pathogenicity tasks classifies non-coding SNVs as benign or pathogenic, and expression QTLs, methylation QTLs, and splicing QTLs classify whether SNVs modify gene expression levels, affect nearby methylation rates or influence alternative splicing, respectively. All VariantBenchmarks use 1,024 bp sequence context. The eQTL task from the Long Range Benchmark (LRB) (8) predicts causal expression quantitative trait loci using 160,000 bp sequence windows and incorporates tissue identity as a conditional input, demonstrating the API’s support for auxiliary features.

##### Zero-shot variant effect tasks

Zero-shot variant effect prediction tasks include BEND, which evaluates disease-associated and expression-associated variants using 512 bp windows; TraitGym (9), which provides Mendelian and complex trait prediction tasks using 4,096 bp contexts; Song-Lab ClinVar (10), which evaluates pathogenicity for clinically annotated SNVs using 5,994 bp windows; and the pathogenic OMIM task from LRB, which evaluates non-coding pathogenic variants using 12,000 bp contexts. We expand this category with an additional task: a standardized ClinVar task aligned with the *vep-eval* framework (11). BRCA1 (12) which evaluates pathogenicity using 8192 bp context window.

##### Zero-shot general indel tasks

Zero-shot general indel tasks include causal eQTL task from LOL-EVE benchmark (13) to evaluate expression modulating insertions and deletions in human promoters. In addition, we introduce a novel ClinVarbased indel pathogenicity task. This task supports insertions and deletions, demonstrating the API’s capability to handle non-aligned sequence inputs, and introducing a novel, open benchmark task for community use.

For evaluation paradigms not addressed by current tasks, new tasks can inherit directly from `BaseGFMTask` and implement the required API methods. All tasks support automatic data acquisition: datasets are downloaded from Hugging-Face, BEND, LRB, or other repositories upon initialization and cached. Variant effect tasks extract sequence contexts from a reference genome (GRCh38) (14), which is downloaded automatically if not present, and sequence windows are centered on variant positions with symmetric truncation applied when user-specified length constraints are more restrictive than task defaults. All concrete task implementations were validated against their original benchmark code-bases or reported results in the corresponding publications, ensuring consistency with published preprocessing pipelines, dataset splits, and evaluation metrics. For additional details on the implemented benchmark tasks, see Appendix, Supplementary Note 5.

## Case Study

To demonstrate GFMBench-API’s versatility, we evaluated five prominent models: DNA-BERT, DNABERT-2, NTv3, Caduceus-Ps, and Evo 2, across our full benchmark suite. This “one-click” workflow decouples model implementation from task logic, enabling standardized comparison of diverse architectures on tasks from supervised variant effect prediction to zero-shot likelihood estimation. Detailed performance metrics are available in the Appendix (Tables 1–5).

**Table 1.** Supervised classification task performance under full fine-tuning. Abbreviations: **MCC** (Matthews Correlation Coefficient), **AUROC** (Area Under the Receiver Operating Characteristic), **AUPRC** (Area Under the Precision-Recall Curve).

| Task | Model | Accuracy | MCC | AUROC | AUPRC |
| --- | --- | --- | --- | --- | --- |
| GUE Transcription Factor | DNA-BERT | 0.8072 | 0.6176 | 0.8963 | 0.8934 |
|  | DNABERT-2 | 0.8212 | 0.6430 | 0.9044 | 0.9023 |
|  | NTv3 | 0.8078 | 0.6167 | 0.8905 | 0.8839 |
|  | Caduceus-Ps | 0.8056 | 0.6140 | 0.8909 | 0.8875 |
| GUE Promoter All | DNA-BERT | 0.9473 | 0.8949 | 0.9903 | 0.9916 |
|  | DNABERT-2 | 0.9306 | 0.8620 | 0.9854 | 0.9861 |
|  | NTv3 | 0.9395 | 0.8791 | 0.9867 | 0.9876 |
|  | Caduceus-Ps | 0.9012 | 0.8024 | 0.9643 | 0.9601 |
| GUE Splice Site | DNA-BERT | 0.9204 | 0.8652 | 0.9824 | 0.9645 |
|  | DNABERT-2 | 0.9226 | 0.8682 | 0.9774 | 0.9568 |
|  | NTv3 | 0.9323 | 0.8841 | 0.9814 | 0.9682 |
|  | Caduceus-Ps | 0.8963 | 0.8207 | 0.9551 | 0.9174 |
| Var Bench Coding Pathogenicity | DNA-BERT | 0.5801 | 0.0858 | 0.5927 | 0.5056 |
|  | DNABERT-2 | 0.6266 | 0.2153 | 0.6973 | 0.5994 |
|  | NTv3 | 0.6221 | 0.2036 | 0.6832 | 0.5793 |
|  | Caduceus-Ps | 0.5745 | 0.0594 | 0.6061 | 0.5068 |
| Var Bench Non Coding Pathogenicity | DNA-BERT | 0.8913 | 0.0720 | 0.6773 | 0.1977 |
|  | DNABERT-2 | 0.8930 | 0.0000 | 0.5050 | 0.1074 |
|  | NTv3 | 0.9805 | 0.8943 | 0.9524 | 0.9314 |
|  | Caduceus-Ps | 0.8930 | 0.0000 | 0.6835 | 0.2407 |
| Var Bench Expression | DNA-BERT | 0.6506 | 0.3336 | 0.6997 | 0.6564 |
|  | DNABERT-2 | 0.6835 | 0.3677 | 0.7490 | 0.7120 |
|  | NTv3 | 0.6745 | 0.3491 | 0.7415 | 0.7059 |
|  | Caduceus-Ps | 0.5998 | 0.2005 | 0.6360 | 0.5921 |
| Var Bench Common Vs Rare | DNA-BERT | 0.5000 | 0.0000 | 0.5005 | 0.5002 |
|  | DNABERT-2 | 0.5000 | 0.0000 | 0.5031 | 0.5020 |
|  | NTv3 | 0.5881 | 0.1803 | 0.6343 | 0.6360 |
|  | Caduceus-Ps | 0.5000 | 0 | 0.5000 | 0.5000 |
| Var Bench meQTL | DNA-BERT | 0.5580 | 0.0708 | 0.6335 | 0.5386 |
|  | DNABERT-2 | 0.5462 | 0.0000 | 0.6186 | 0.5147 |
|  | NTv3 | 0.5729 | 0.1185 | 0.6556 | 0.5783 |
|  | Caduceus-Ps | 0.5462 | 0.0078 | 0.5648 | 0.4884 |
| Var Bench sQTL | DNA-BERT | 0.4949 | 0.0102 | 0.5205 | 0.5263 |
|  | DNABERT-2 | 0.5049 | 0.0589 | 0.5485 | 0.5581 |
|  | NTv3 | 0.5022 | 0.0402 | 0.5404 | 0.5463 |
|  | Caduceus-Ps | 0.5549 | 0.1062 | 0.5681 | 0.5696 |
| LRB Variant Effect Causal eQTL | DNA-BERT | 0.6012 | 0.2317 | 0.6328 | 0.6573 |
|  | DNABERT-2 | 0.6291 | 0.2853 | 0.6780 | 0.6850 |
|  | NTv3 | 0.6248 | 0.2798 | 0.6804 | 0.7019 |
|  | Caduceus-Ps | 0.6286 | 0.2838 | 0.6792 | 0.6856 |

## Conclusion

As genomic foundation models grow in complexity, GFMBench-API addresses the critical need for rigorous evaluation by replacing fragmented, ad-hoc scripts with a standardized engineering protocol. By decoupling biological task logic from model architecture, this unified interface enables a “plug-and-play” ecosystem for testing new models against established benchmarks. GFMBench-API lowers the barrier to entry for genomic AI development and ensures that performance claims are validated through a consistent, rigorous framework fostering a more systematic understanding of progress in the field. While the current framework is primarily tailored to the haploid genomic domain, future extensions will focus on broadening task coverage and incorporating more complex scenarios, including diploid and other multi-context sequence analyses, to further enhance its applicability and scope.

## Software Availability

GFMBench-API is implemented in Python and and is freely available for academic use. Code will be publicly released upon acceptance.

## Data availability

The data supporting this study are accessible online and can also be obtained by contacting the corresponding author.

## Author contributions statement

A.L., E.D., A.B., Y.S. and N.D. conceived the study, designed and developed the methodology, performed the experiments, and analysed the results. R.K., G.L., O.N., D.O., T.Z., D.D., and G.R. contributed to method development, performed experiments, data analysis, and critical review of the results. N.B., S.L., S.O., D.H., M.W., and A.C. contributed to manuscript review and provided scientific feedback. Y.S. and N.D. supervised the project, guided the research direction. All authors reviewed and approved the final manuscript.

## Appendix

Our appendix provides additional implementation details.

### Supplementary Note 1: Comparative Study of GFMs Benchmarks

Several recent studies have advanced the evaluation of Genomic Foundation Models (GFMs) from complementary perspectives. DNA Foundation Benchmarks (15) present a systematic comparison of DNA foundation models using frozen zero-shot embeddings across a wide range of genomic tasks, highlighting the impact of embedding aggregation strategies and task dependency on downstream performance. GenBench (16) focuses on benchmarking short- and long range genomic tasks, providing insights into how different architectural families exhibit distinct performance characteristics across genomic scale. OmniGenBench (17) further contributes an automated benchmarking platform that integrates large-scale datasets and standardized execution pipelines to improve reproducibility and accessibility of GFM evaluation. Compared to these efforts, GFMBench-API does not introduce new benchmark datasets or evaluation metrics, but instead proposes a standardized application programming interface that unifies task definitions, model inference outputs, and metric computation under a common abstraction. Compared to benchmark frameworks (18), (17), which couple task logic to specific model implementations and require model adaptations per task, GFMBench-API adopts a decoupled design in which models expose a small set of generic inference capabilities that can be reused across heterogeneous task types and metrics. This property is particularly important in genomics, given the diversity of supervised, zero-shot, and comparative reference / variant settings. Additionally, GFMsBench-API is intentionally domain-centered, focusing exclusively on DNA-related tasks to evaluate GFMs with the aim of improving long-term sustainability and usability.

### Supplementary Note 2: Evaluation Metrics

This section describes the evaluation metrics used to benchmark genomic foundation models across supervised and zero-shot tasks.

#### Classification Accuracy

This metric measures the proportion of correctly classified samples. For each input sequence, the predicted label is obtained by taking the argmax over the model’s output probability distribution across classes. Accuracy is computed as the number of correct predictions divided by the total number of samples. While straightforward and interpretable, this metric can be sensitive to class imbalance and is therefore reported alongside more robust measures.

#### Classification MCC

Quantifies classification performance using the Matthews Correlation Coefficient (MCC), which accounts for true positives, true negatives, false positives, and false negatives. Predictions are obtained via argmax over class probabilities. MCC ranges from − 1 to 1, where 1 indicates perfect prediction, 0 corresponds to random performance, and − 1 indicates complete disagreement. This metric is particularly informative for imbalanced datasets.

#### Classification AUROC

This metric evaluates the model’s ability to discriminate between classes across all possible decision thresholds. For binary classification, AUROC is computed using the predicted probability of the positive class. For multi-class problems, a macro-averaged one-vs-rest (OVR) AUROC is reported. An AUROC of 0.5 indicates random performance, while 1.0 denotes perfect discrimination.

#### Classification AUPRC

This metric summarizes the trade-off between precision and recall across decision thresholds. For binary classification, AUPRC is computed using the predicted probability of the positive class, while for multi-class settings a macro-averaged precision–recall score is used. AUPRC is particularly informative for imbalanced datasets, where baseline performance depends on class prevalence.

#### Mean Probs LLR AUROC / AUPRC

This zero-shot metric compares the log-likelihood of a variant sequence to that of its corresponding reference sequence under the model. For autoregressive models, token-level log-probabilities are obtained from the next-token prediction head, and the log-probabilities of the observed tokens are averaged to compute normalized sequence log-likelihoods. The log-likelihood ratio (LLR) is then defined as LL_var_ − LL_ref_. Because pathogenic variants are typically assigned lower likelihoods than their reference counterparts, the negative LLR is used as the prediction score. Performance is evaluated using both AUROC and AUPRC.

#### Seq Embeddings Cosine Sim AUROC / AUPRC

This metric measures the cosine similarity between sequence-level representative embeddings of the variant and reference sequences. Pathogenic variants are expected to induce larger semantic changes, resulting in lower cosine similarity; therefore, the negative cosine similarity is used as the prediction score and evaluated using AUROC and AUPRC.

#### Seq Embeddings L2 AUROC / AUPRC

This metric computes the Euclidean (L2) distance between sequence-level representative embeddings of the variant and reference sequences. This metric captures the magnitude of the embedding shift induced by the variant. Larger distances are hypothesized to correspond to more disruptive variants. The L2 distance is used directly as the prediction score and evaluated using AUROC and AUPRC.

#### SNV Variant Effect Cosine Sim. AUROC / AUPRC

This is an, SNV-specific metric that focuses on local embedding changes at the mutation site. Cosine similarity is computed between token-level embeddings at the SNV position in the variant and reference sequences. Pathogenic variants are expected to cause larger local embedding deviations, leading to lower cosine similarity. The negative cosine similarity is evaluated using AUROC and AUPRC.

#### SNV Variant Effect Pred Masked LLR AUROC / AUPRC

This metric assesses variant impact using masked language modeling predictions at the SNV position. For masked language models, the input sequence is processed with the SNV position masked, and the model predicts a probability distribution over nucleotides at the masked position using the masked language modeling head employed during pretraining. The probabilities assigned to the reference and variant nucleotides are extracted, and the log-likelihood ratio is computed as log *P* (variant) − log *P* (reference). Since pathogenic variants are generally assigned lower probability than the reference nucleotide, the negative LLR is used as the prediction score and evaluated using AUROC and AUPRC.

### Supplementary Note 3: Model Inference Methods

This section documents the collection of standardized inference methods that models may implement to be compatible with the GFMBench-API. These methods define the interface between user-provided models and benchmark tasks, enabling task-agnostic evaluation across diverse model architectures, training objectives, and tokenization strategies. Models may implement any subset of these methods, and tasks automatically invoke only those required to compute their associated evaluation metrics. This section documents the collection of standardized inference methods that models may implement to be compatible with the GFMBench-API. These methods define the interface between user-provided models and benchmark tasks, enabling task-agnostic evaluation across diverse model architectures, training objectives, and tokenization strategies. Across all inference methods, input DNA sequences are represented as strings over the alphabet A, C, G, T, representing the four canonical nucleotides, with the additional symbol N denoting unknown or ambiguous nucleotides and the symbol P denoting padding. Models may implement any subset of these methods, and tasks automatically invoke only those required to compute their associated evaluation metrics.

#### Single-Sequence Classification Inference

This method maps individual DNA sequences to probability distributions over a fixed set of categorical labels. It is used by supervised single-sequence classification tasks, such as promoter prediction or splice site classification. The output probabilities are expected to sum to one across labels and are used to compute standard classification metrics, including accuracy, MCC, AUROC, and AUPRC. Optional conditional inputs may be provided to support tasks that incorporate auxiliary metadata.

#### Variant-Reference Pair Classification Inference

This method computes classification probabilities for paired variant and reference sequences in supervised variant effect prediction tasks. It enables models to jointly reason over the reference and alternate alleles to predict functional or pathogenic labels. The specific strategy for combining variant and reference information is model-dependent and may include Siamese architectures, sequence concatenation, or difference-based encodings.

#### Sequence-to-Sequence Inference

This method operates on a batch of DNA sequences and produces sequence-level and/or token-level outputs. Given a DNA sequence, the method may return up to three optional outputs within a single inference call, depending on the model architecture and implemented inference capabilities.

The first output consists of token-level embeddings for each sequence position, representing the hidden states produced by the model at each token. The second output consists of per-position token probabilities corresponding to next-token prediction, where each probability represents the likelihood assigned by the model to the observed token at that position. The third output is a single representative embedding for each sequence, intended to summarize the entire sequence. The construction of this representative embedding is model-defined and may correspond to a special classification token (e.g., CLS), an end-of-sequence token, or an aggregation such as mean pooling over token embeddings.

These outputs are exposed through a unified inference interface to avoid redundant forward passes through the model. This design enables efficient evaluation by allowing multiple metrics to be computed from a single model invocation. At least one of these outputs must be provided for meaningful evaluations, and the benchmark framework automatically computes only those metrics compatible with the outputs returned by the model.

#### Masked Token Prediction Inference

This method performs masked language model inference at a specified variant position. The input sequence is processed with the nucleotide at the variant position masked, and the model predicts the probabilities of both the reference and alternate nucleotides at that position, accounting for the model’s tokenization scheme. This inference mode is relevant only for models trained with masked language modeling objectives and is not required for autoregressive models.

#### Sequence Position Mapping

This method maps nucleotide positions in the input DNA sequence to corresponding positions in the model’s output space. It is required for position-specific metrics in zero-shot SNV tasks, where tokenization schemes (e.g., k-mer or BPE tokenization) may cause misalignment between input nucleotides and output embeddings or probabilities. The method enables consistent positional alignment across models with different tokenization strategies.

### Supplementary Note 4: Conditional Inputs and Metadata Support

GFMBench-API supports optional conditional inputs that provide auxiliary information alongside DNA sequences during model inference. Conditional inputs enable context-aware and multi-modal evaluation settings, such as tissue-specific variant effect prediction, while remaining fully compatible with tasks and models.

At the dataset level, each sample may include an additional conditional input vector. For single-sequence tasks, samples take the form (sequence, label, conditional input), while variant effect tasks use (variant sequence, reference sequence, label, conditional input). Conditional inputs are represented as numeric vectors and, when batched, are passed to models as arrays of shape (batch size, number of features).

All model inference methods defined by the API accept an optional conditional input argument. Models may ignore this input, incorporate it as an additional conditioning signal, or use it in an architecture-specific manner.

Tasks that provide conditional inputs expose a metadata schema describing each auxiliary feature through a dedicated task-level interface. This schema specifies the name, data type, and expected value range of each metadata feature, and the ordering of features corresponds directly to the indices of the conditional input vector. The metadata schema is included in the task attributes, allowing benchmark orchestration code and model implementations to inspect metadata requirements programmatically. Tasks that do not provide such metadata remain fully operational, as the corresponding metadata argument is returned as an empty indicator. Furthermore, for datasets where some samples lack metadata entries, this decoupling of model development and inference from task design allows developers to flexibly manage missing metadata samples after these empty indicators are supplied for each affected input vector. This ensures that both task-level and sample-level metadata gaps are handled seamlessly within the unified framework.

This design enables flexible integration of auxiliary biological context while preserving model agnosticism, efficient batched inference, and consistent evaluation across tasks. Although only a subset of current benchmark tasks utilize conditional inputs, the framework is designed to support a wide range of metadata-rich genomic evaluation scenarios.

### Supplementary Note 5: Description of Implemented Benchmark Tasks

This section provides detailed descriptions of all benchmark tasks implemented using the proposed API. For each task, we describe the task objective and data formulation, followed by differences from the original benchmark implementation when applicable and the methodology used to reproduce reported results.

#### A. Supervised Single-Sequence Classification Tasks

All supervised single-sequence tasks operate on fixed-length DNA sequences paired with categorical labels and use predefined train and test data splits (and may also have a validation set). Evaluation is performed using standard classification metrics, including accuracy, Matthews correlation coefficient (MCC), AUROC, and AUPRC.

##### GUE Promoter Prediction

This task from the GUE benchmark evaluates promoter prediction as a binary classification problem using 300 bp DNA sequences labeled as promoter or non-promoter. Results are reproduced by fine-tuning models on the training set for a fixed number of epochs and evaluating on the test set, following the protocol reported in the GUE benchmark.

##### GUE Splice Site Prediction

This GUE benchmark task formulates splice site recognition as a three-class classification problem, labeling 400 bp sequence windows as donor, acceptor, or non-splice. Results are reproduced by end-to-end fine-tuning and evaluation on the predefined test split using MCC as the primary metric.

##### GUE Transcription Factor Binding Site Prediction

This task from the GUE benchmark predicts whether a 100 bp DNA sequence corresponds to a transcription factor binding site. Sequences from multiple ChIP-seq experiments are aggregated into a single binary classification dataset. Results are reproduced by fine-tuning models on the provided training data and evaluating on the test split.

#### A. Supervised Variant Effect Prediction Tasks

Supervised variant effect prediction tasks operate on paired reference and alternate sequences and predict binary functional labels. Sequence contexts are centered on the variant position. Similarly to the single-Sequence classification tasks, the input sequences are paired with categorical labels and use predefined data splits, and the classification evaluation metrics.

##### VariantBenchmarks Coding Pathogenicity

This task from the VariantBenchmarks suite classifies single-nucleotide variants in protein-coding regions as benign or pathogenic using 1,024 bp sequence contexts. Chromosomes 21, 22, and X are reserved for testing. Unlike the original benchmark, which reports cross-validated results, a fixed test fold is used. Results are reproduced using linear probing, where aggregated reference and alternate embeddings are used to train a logistic regression classifier.

##### VariantBenchmarks Non-Coding Pathogenicity

This VariantBenchmarks task evaluates pathogenicity prediction for non-coding variants using the same sequence length and chromosome-based test split as the coding task. A fixed-fold evaluation strategy is also used instead of cross-validation, and the results are reproduced using linear probing with logistic regression on aggregated embeddings.

##### VariantBenchmarks Common vs. Rare Variant Classification

This task evaluates whether models capture the genomic characteristic of common variants, defined by a minor allele frequency less than 0.05, compared to synthetic rare controls. The results are reproduced using linear probing on aggregated reference and alternate embeddings with logistic regression.

##### VariantBenchmarks Expression, meQTL, and sQTL Prediction

These VariantBenchmarks tasks evaluate whether a variant affects gene expression levels, DNA methylation rates, or alternative splicing, respectively. All tasks use 1,024 bp sequence contexts and chromosome-based test splits. The results are reproduced using a shared linear probing setup with logistic regression.

##### Long Range Benchmark Causal eQTL Prediction

This task from the Long Range Benchmark predicts whether a variant causally influences gene expression using sequence windows capable of spanning up to 160,000 bp, and incorporates tissue identity as an auxiliary conditional input. The results are reproduced by applying the aggregated reference and alternate embeddings to a projection layer, followed by fine-tuning and evaluating the combined model.

#### C. Zero-Shot SNV Variant Effect Prediction Tasks

Zero-shot SNV tasks evaluate pretrained models without task-specific fine-tuning. These tasks compare reference and variant DNA sequences using likelihood-based scores, sequence-level embeddings, and token-level embedding–based measures. The variant sequences correspond to single-nucleotide variants (SNVs); therefore, the reference and variant sequences are required to have identical lengths.

##### BEND Variant Effect Prediction (Disease, Expression)

This task from the BEND benchmark evaluates disease-associated and expression-associated non-coding variants using 512 bp windows centered on the variant position. Results are reproduced by running zero-shot evaluation on the full dataset.

##### Song Lab ClinVar Pathogenicity

This task evaluates ClinVar-annotated single-nucleotide variants using 5,994 bp sequence contexts as inputs for the pathogenisity classification task. Results are reproduced by validating against the original Song Lab evaluation code on the masked LLR–based metrics.

##### VEP-Eval ClinVar Pathogenicity

This task aligns ClinVar evaluation with the VEP-Eval framework. Compared to the reference implementation, samples with missing predictions are removed prior to evaluation. Results are reproduced using embedding-distance-based scoring.

##### BRCA1 Pathogenicity Prediction

This task evaluates the ability of models to predict the functional impact of single nucleotide variants (SNVs) in the *BRCA*1 gene. The dataset consists of 3,893 variants categorized as loss-of-function (LOF), intermediate (INT), or functional (FUNC) based on experimentally measured DNA-repair activity. Results are reproduced by calculating zero-shot likelihood scores for reference and alternate alleles and assessing the model’s classification accuracy via AUROC and correlation with experimental function scores.

##### TraitGym

These tasks from TraitGym evaluate zero-shot prediction of Mendelian and complex trait–associated variants using 4,096 bp sequence contexts. Compared to the original evaluation, reverse-complement averaging and repeated bootstrapping are omitted. Results are reproduced using embedding-distance-based scores.

##### Long Range Benchmark Pathogenic OMIM Prediction

This task from the Long Range Benchmark evaluates non-coding pathogenic variants associated with Mendelian diseases using sequence contexts capable of spanning up to 160,000 bp. The results are reproduced by zero-shot evaluation on a filtered subset of SNVs.

#### D. Zero-Shot General Indel Tasks

Zero-shot General Indel tasks evaluate pretrained models without task-specific fine-tuning. Reference and alternate sequences are compared using likelihood-based and embedding-based scores. The alternate sequences may correspond to insertion or deletion (indel) variants and therefore can differ in length from the reference sequence.

##### LOL-EVE Causal eQTL

This task from the LOL-EVE benchmark evaluates causal expression modulating variants (insertions and deletions) in human promoters. Results are reproduced using zero-shot unmasked reconstruction scoring: per-position probabilities for the observed nucleotide tokens are obtained from a single unmasked forward pass and aggregated as the mean log-probability over valid tokens. Variant impact is scored as the absolute ALT–REF log-likelihood ratio. The evaluation follows a replicated process using posterior inclusion probability (PIP) and sequence slippage thresholds, validated across the range of described experimental criteria.

##### ClinVar Indel Pathogenicity

This task evaluates insertions and deletions from ClinVar for pathogenicity in a zero-shot setting using non-aligned reference and alternate sequences. The task supports general indels and excludes position-specific SNV metrics. This task introduces a new open benchmark. Results are reproduced using sequence-level likelihood- and embedding-based scoring.

### Supplementary Note 6: Evaluation of GFMs using GFMBench-API

This section outlines the evaluation methodology using GFMBench-API to benchmark performance in various genomic foundation models. We report results for DNA-BERT (1), DNABERT-2 (3), Nucleotide Transformer v3 (NTv3) (5), Caduceus-Ps (19), and Evo2 (20). The comprehensive performance metrics for the supervised classification, zero-shot likelihood, and zero-shot embedding tasks are detailed in Tables 1, 2, 3, 4, and, 5 respectively.

**Table 2.** Supervised classification task performance for **Evo2** under linear probing. Abbreviations: **MCC** (Matthews Correlation Coefficient), **AUROC** (Area Under the Receiver Operating Characteristic), **AUPRC** (Area Under the Precision-Recall Curve).

| Task | Accuracy | MCC | AUROC | AUPRC |
| --- | --- | --- | --- | --- |
| GUE Transcription Factor | 0.5850 | 0.1765 | 0.6305 | 0.6016 |
| GUE Promoter All | 0.7539 | 0.5102 | 0.8149 | 0.7781 |
| GUE Splice Site | 0.5649 | 0.0384 | 0.6571 | 0.4601 |
| Var Bench Coding Pathogenicity | 0.5801 | 0.0941 | 0.6585 | 0.5677 |
| Var Bench Non Coding Pathogenicity | 0.8980 | 0.2045 | 0.6590 | 0.4155 |
| Var Bench Expression | 0.5949 | 0.1974 | 0.6278 | 0.5893 |
| Var Bench Common Vs Rare | 0.5152 | 0.0390 | 0.5245 | 0.5180 |
| Var Bench meQTL | 0.5393 | -0.0031 | 0.5665 | 0.4847 |
| LRB Variant Effect Causal eQTL | 0.6152 | 0.2302 | 0.6618 | 0.6411 |

**Table 3.** Zero-Shot Embedding Metrics. Abbreviations: **Cos** (Sequence Embeddings Cosine Similarity), **L2** (Sequence Embeddings L2 Distance), **SNV Cos** (SNV Variant Effect Cosine Similarity), **NA** (Indicates that SNV metrics are not applicable to indel tasks.).

| Task | Model | Cos AUROC | Cos AUPRC | L2 AUROC | L2 AUPRC | SNV Cos AUROC | SNV Cos AUPRC |
| --- | --- | --- | --- | --- | --- | --- | --- |
| Bend Variant Effects Disease | DNA-BERT | 0.5086 | 0.0730 | 0.5088 | 0.0731 | 0.5345 | 0.0731 |
|  | DNABERT-2 | 0.6962 | 0.1519 | 0.6928 | 0.1489 | 0.5115 | 0.0751 |
|  | Evo2 | 0.8723 | 0.4738 | 0.9116 | 0.5955 | 0.6149 | 0.0944 |
|  | NTv3 | 0.7128 | 0.1471 | 0.7196 | 0.1522 | 0.7681 | 0.2749 |
|  | Caduceus-Ps | 0.7724 | 0.3782 | 0.7798 | 0.3788 | 0.5446 | 0.1071 |
| Bend Variant Effects Expression | DNA-BERT | 0.5305 | 0.0812 | 0.5311 | 0.0814 | 0.5976 | 0.0935 |
|  | DNABERT-2 | 0.5478 | 0.0848 | 0.5515 | 0.0853 | 0.4871 | 0.0732 |
|  | Evo2 | 0.5820 | 0.0904 | 0.5732 | 0.0897 | 0.5569 | 0.0887 |
|  | NTv3 | 0.5140 | 0.0789 | 0.5321 | 0.0818 | 0.4472 | 0.0640 |
|  | Caduceus-Ps | 0.4729 | 0.0676 | 0.4794 | 0.0692 | 0.5371 | 0.0775 |
| Songlab ClinVar | DNA-BERT | 0.5101 | 0.5499 | 0.5100 | 0.5500 | 0.5392 | 0.5668 |
|  | DNABERT-2 | 0.4963 | 0.5472 | 0.4955 | 0.5471 | 0.5008 | 0.5523 |
|  | Evo2 | 0.7220 | 0.7341 | 0.8445 | 0.8654 | 0.5790 | 0.5761 |
|  | NTv3 | 0.5085 | 0.5524 | 0.5157 | 0.5591 | 0.5554 | 0.5925 |
|  | Caduceus-Ps | 0.9333 | 0.9958 | 0.8667 | 0.9914 | 0.8667 | 0.9914 |
| TraitGym Complex | DNA-BERT | 0.5091 | 0.1039 | 0.5095 | 0.1038 | 0.5387 | 0.1107 |
|  | DNABERT-2 | 0.5296 | 0.1089 | 0.5358 | 0.1115 | 0.4983 | 0.0986 |
|  | Evo2 | 0.5274 | 0.1077 | 0.5236 | 0.1068 | 0.5228 | 0.1062 |
|  | NTv3 | 0.5068 | 0.1032 | 0.5184 | 0.1058 | 0.4998 | 0.1018 |
|  | Caduceus-Ps | 0.4835 | 0.0946 | 0.4967 | 0.0974 | 0.5177 | 0.1034 |
| TraitGym Mendelian | DNA-BERT | 0.5291 | 0.1061 | 0.5301 | 0.1063 | 0.5970 | 0.1291 |
|  | DNABERT-2 | 0.5939 | 0.1473 | 0.6051 | 0.1533 | 0.4629 | 0.0890 |
|  | Evo2 | 0.6176 | 0.1641 | 0.6263 | 0.1702 | 0.5480 | 0.1211 |
|  | NTv3 | 0.5370 | 0.1030 | 0.5531 | 0.1072 | 0.4813 | 0.0933 |
|  | Caduceus-Ps | 0.4629 | 0.0879 | 0.4775 | 0.0922 | 0.5527 | 0.1273 |
| LRB Variant Effect OMIM | DNA-BERT | 0.5140 | 0.0021 | 0.5142 | 0.0021 | 0.5625 | 0.0027 |
|  | DNABERT-2 | 0.5123 | 0.0021 | 0.5308 | 0.0022 | 0.5052 | 0.0019 |
|  | Evo2 | 0.5008 | 0.0020 | 0.5016 | 0.0021 | 0.4980 | 0.0020 |
|  | NTv3 | 0.4798 | 0.0018 | 0.5001 | 0.0019 | 0.4739 | 0.0018 |
|  | Caduceus-Ps | 0.4863 | 0.0019 | 0.4901 | 0.0019 | 0.5240 | 0.0022 |
| BRCA1 | DNA-BERT | 0.5137 | 0.2158 | 0.5141 | 0.2159 | 0.5271 | 0.2267 |
|  | DNABERT-2 | 0.4868 | 0.2141 | 0.4760 | 0.1990 | 0.4903 | 0.2065 |
|  | Evo2 | 0.7468 | 0.5401 | 0.7752 | 0.5745 | 0.4668 | 0.2041 |
|  | NTv3 | 0.5571 | 0.2458 | 0.5620 | 0.2498 | 0.6244 | 0.3029 |
|  | Caduceus-Ps | 0.5553 | 0.2910 | 0.5631 | 0.2986 | 0.4690 | 0.2025 |
| VEP-eval | DNA-BERT | 0.5113 | 0.3101 | 0.5114 | 0.3104 | 0.5961 | 0.3665 |
|  | DNABERT-2 | 0.4735 | 0.2912 | 0.4731 | 0.2915 | 0.4848 | 0.2929 |
|  | NTv3 | 0.6840 | 0.4719 | 0.6912 | 0.4790 | 0.5845 | 0.3829 |
|  | Caduceus-Ps | 0.7023 | 0.5561 | 0.7073 | 0.5627 | 0.5109 | 0.3633 |
| ClinVar Indel Pathogenicity | DNA-BERT | 0.5008 | 0.8533 | 0.5011 | 0.8538 | NA | NA |
|  | DNABERT-2 | 0.5080 | 0.8598 | 0.5082 | 0.8616 | NA | NA |
|  | Evo2 | 0.9458 | 0.9885 | 0.9666 | 0.9926 | NA | NA |
|  | NTv3 | 0.6416 | 0.9437 | 0.6475 | 0.9456 | NA | NA |
|  | Caduceus-Ps | 0.7080 | 0.9581 | 0.7100 | 0.9585 | NA | NA |
| LOL-EVE | DNA-BERT | 0.4477 | 0.0676 | 0.4490 | 0.0681 | NA | NA |
|  | DNABERT-2 | 0.6014 | 0.1050 | 0.5787 | 0.0947 | NA | NA |
|  | Evo2 | 0.5313 | 0.0845 | 0.5435 | 0.0938 | NA | NA |
|  | NTv3 | 0.4948 | 0.0867 | 0.5031 | 0.0903 | NA | NA |
|  | Caduceus-Ps | 0.5242 | 0.0861 | 0.5216 | 0.0863 | NA | NA |

**Table 4.** Zero-Shot Likelihood: **Masked Language Models.** Performance measured via Masked Log-Likelihood Ratios (LLR).

| Task | Model | Msk LLR ROC | Msk LLR PRC |
| --- | --- | --- | --- |
| Bend Variant Effects Disease | DNA-BERT | 0.4320 | 0.0593 |
|  | DNABERT-2 | 0.5044 | 0.0706 |
|  | NTv3 | 0.6646 | 0.1584 |
|  | Caduceus-Ps | 0.6598 | 0.1397 |
| Bend Variant Effects Expression | DNA-BERT | 0.4787 | 0.0681 |
|  | DNABERT-2 | 0.5004 | 0.0732 |
|  | NTv3 | 0.4871 | 0.0682 |
|  | Caduceus-Ps | 0.4886 | 0.0679 |
| Songlab ClinVar | DNA-BERT | 0.4515 | 0.5031 |
|  | DNABERT-2 | 0.5189 | 0.5530 |
|  | NTv3 | 0.5496 | 0.5826 |
|  | Caduceus-Ps | 0.5222 | 0.5459 |
| TraitGym Complex | DNA-BERT | 0.4992 | 0.0981 |
|  | DNABERT-2 | 0.4927 | 0.0977 |
|  | NTv3 | 0.4899 | 0.0941 |
|  | Caduceus-Ps | 0.4827 | 0.0923 |
| TraitGym Mendelian | DNA-BERT | 0.4691 | 0.0905 |
|  | DNABERT-2 | 0.4819 | 0.0949 |
|  | NTv3 | 0.5089 | 0.0957 |
|  | Caduceus-Ps | 0.5046 | 0.0929 |
| LRB Variant Effect OMIM | DNA-BERT | 0.5225 | 0.0021 |
|  | DNABERT-2 | 0.4771 | 0.0019 |
|  | NTv3 | 0.4634 | 0.0017 |
|  | Caduceus-Ps | 0.4105 | 0.0016 |
| VEP-eval | DNA-BERT | 0.4243 | 0.2620 |
|  | DNABERT-2 | 0.5216 | 0.3117 |
|  | NTv3 | 0.7124 | 0.5728 |
|  | Caduceus-Ps | 0.6616 | 0.4777 |
| BRCA1 | DNA-BERT | 0.4336 | 0.1789 |
|  | DNABERT-2 | 0.5582 | 0.2386 |
|  | NTv3 | 0.4932 | 0.2214 |
|  | Caduceus-Ps | 0.4832 | 0.2180 |

**Table 5.** Zero-Shot Likelihood: **Evo2** (Autoreggresive). Performance is quantified by Log-Likelihood Ratios (LLR), computed over mean predicted token probabilities.

| Task | Mean LLR ROC | Mean LLR PRC |
| --- | --- | --- |
| Bend Variant Effects Disease | 0.9194 | 0.7422 |
| Bend Variant Effects Expression | 0.4885 | 0.0685 |
| Songlab ClinVar | 0.9215 | 0.9327 |
| TraitGym Complex | 0.4920 | 0.0939 |
| TraitGym Mendelian | 0.6041 | 0.1607 |
| LRB Variant Effect OMIM | 0.5080 | 0.0024 |
| BRCA1 | 0.7412 | 0.5328 |
| ClinVar Indel Pathogenicity | 0.6270 | 0.8919 |
| LOL-EVE | 0.5477 | 0.1087 |

#### A. Model Sourcing and Implementation

All models were sourced from Hugging Face (21), except for Evo2, which was implemented using the NVIDIA BioNeMo framework (22). DNA-BERT is a 110M-parameter model, while DNABERT-2 contains approximately 117M parameters. NTv3 is a transformer with approximately 8M parameters, and Caduceus-Ps comprises around 7M parameters. Evo2 is a 1B-parameter model.

#### B. Fine-Tuning and Training Protocols

For the supervised learning tasks presented in Tables 1 and 2, different training strategies were used:

- Full Fine-Tuning: DNA-BERT, DNABERT-2, NTv3, and Caduceus-Ps underwent full fine-tuning for 3 epochs to optimize their weights for specific downstream tasks.
- Linear Probing: In contrast, Evo 2 was evaluated using a linear probing approach. The pre-trained model weights were frozen, and only a linear classification head was trained, ensuring the core model parameters remained static.

#### C. Methodology for Supervised Variant Effect Prediction

For supervised variant effect tasks, a dual-inference strategy was utilized across all models. Both the reference and variant sequences were independently inferred to generate their respective sequence embeddings. These embeddings were concatenated and projected through a final classification layer to produce class logits

#### D. Zero-Shot Evaluation

Beyond supervised fine-tuning, models were also assessed in a zero-shot setting to evaluate their intrinsic capabilities without task-specific training. These evaluations focus on embedding-based similarity metrics (Table 3), and likelihood-based metrics (Tables 4 and 5), providing insight into the models’ generalized understanding of genomic sequences

#### E. Hardware Configuration

DNA-BERT and DNABERT-2 tasks (both zero-shot and fine-tuning) were executed on a single NVIDIA A10G GPU with 24GB VRAM. NTv3 and Caduceus-Ps tasks (both zero-shot and fine-tuning) were executed on a single NVIDIA A100 GPU with 80GB VRAM. Evo2 evaluations, including both zero-shot and linear probing runs, were conducted at half precision (FP16) on a single NVIDIA H100 GPU with 80GB VRAM

## Bibliography

1. Yanrong Ji, Zhihan Zhou, Han Liu, and Ramana V Davuluri. Dnabert: pre-trained bidirectional encoder representations from transformers model for dna-language in genome. Bioinformatics, 37(15):2112–2120, 2021.

2. Eric Nguyen, Michael Poli, Matthew G Durrant, Brian Kang, Dhruva Katrekar, David B Li, Liam J Bartie, Armin W Thomas, Samuel H King, Garyk Brixi, et al. Sequence modeling and design from molecular to genome scale with evo. Science, 386(6723):eado9336, 2024.

3. Zhihan Zhou, Yanrong Ji, Weijian Li, Pratik Dutta, Ramana Davuluri, and Han Liu. Dnabert-2: Efficient foundation model and benchmark for multi-species genomes. In International Conference on Learning Representations, volume 2024, pages 41642–41665, 2024.

4. Frederikke Marin, Felix Teufel, Marc Horlacher, Dennis Madsen, Dennis Pultz, Ole Winther, and Wouter Boomsma. Bend: Benchmarking dna language models on biologically meaningful tasks. In International Conference on Learning Representations, volume 2024, pages 15246–15281, 2024.

5. Hugo Dalla-Torre, Liam Gonzalez, Javier Mendoza-Revilla, Nicolas Lopez Carranza, Adam Henryk Grzywaczewski, Francesco Oteri, Christian Dallago, Evan Trop, Bernardo P De Almeida, Hassan Sirelkhatim, et al. Nucleotide transformer: building and evaluating robust foundation models for human genomics. Nature methods, 22(2):287–297, 2025.

6. Melissa J Landrum, Jennifer M Lee, George R Riley, Wonhee Jang, Wendy S Rubinstein, Deanna M Church, and Donna R Maglott. Clinvar: public archive of relationships among sequence variation and human phenotype. Nucleic acids research, 42(D1):D980–D985, 2014.

7. Aleksandr Medvedev, Karthik Viswanathan, Praveenkumar Kanithi, Kirill Vishniakov, Prateek Munjal, Clement Christophe, Marco AF Pimentel, Ronnie Rajan, and Shadab Khan. Biotoken and biofm–biologically-informed tokenization enables accurate and efficient genomic foundation models. bioRxiv, pages 2025–03, 2025.

8. Chia Hsiang Kao, Evan Trop, McKinley Polen, Yair Schiff, Bernardo P de Almeida, Aaron Gokaslan, Thomas Pierrot, and Volodymyr Kuleshov. Advancing dna language models: The genomics long-range benchmark. In ICLR 2024 Workshop on Machine Learning for Genomics Explorations, 2024.

9. Gonzalo Benegas, Gökcen Eraslan, and Yun S Song. Benchmarking dna sequence models for causal regulatory variant prediction in human genetics. bioRxiv, 2025.

10. Gonzalo Benegas, Carlos Albors, Alan J Aw, Chengzhong Ye, and Yun S Song. A dna language model based on multispecies alignment predicts the effects of genome-wide variants. Nature Biotechnology, 43(12):1960–1965, 2025.

11. Baiyu Lu, Xueshen Liu, Po-Yu Lin, and Nadav Brandes. Genomic heterogeneity inflates the performance of variant pathogenicity predictions. bioRxiv, pages 2025–09, 2025.

12. Gregory M Findlay, Riza M Daza, Beth Martin, Melissa D Zhang, Anh P Leith, Molly Gasperini, Joseph D Janizek, Xingfan Huang, Lea M Starita, and Jay Shendure. Accurate classification of brca1 variants with saturation genome editing. Nature, 562(7726):217–222, 2018.

13. 1. Courtney A Shearer, Rose Orenbuch, Felix Teufel, Christian J Steinmetz, Daniel Ritter, Erik Xie, Artem Gazizov, Aviv Spinner, Jonathan Frazer, Mafalda Dias, et al. A genomic language model for zero-shot prediction of promoter variant effects. bioRxiv, pages 2024–11, 2024.

14. Valerie A Schneider, Tina Graves-Lindsay, Kerstin Howe, Nathan Bouk, Hsiu-Chuan Chen, Paul A Kitts, Terence D Murphy, Kim D Pruitt, Françoise Thibaud-Nissen, Derek Albracht, et al. Evaluation of grch38 and de novo haploid genome assemblies demonstrates the enduring quality of the reference assembly. Genome research, 27(5):849–864, 2017.

15. Haonan Feng, Lang Wu, Bingxin Zhao, Chad Huff, Jianjun Zhang, Jia Wu, Lifeng Lin, Peng Wei, and Chong Wu. Benchmarking dna foundation models for genomic and genetic tasks. Nature communications, 16(1):10780, 2025.

16. Zicheng Liu, Jiahui Li, Siyuan Li, Zelin Zang, Cheng Tan, Yufei Huang, Yajing Bai, and Stan Z Li. Genbench: A benchmarking suite for systematic evaluation of genomic foundation models. arXiv preprint arXiv:2406.01627, 2024.

17. Heng Yang, Jack Cole, and Ke Li. Omnigenbench: Automating large-scale in-silico bench-marking for genomic foundation models. arXiv preprint arXiv:2410.01784, 2024.

18. Elizabeth Fahsbender, Alma Andersson, Jeremy Ash, Polina Binder, Daniel Burkhardt, Benjamin Chang, Georg K Gerber, Anthony Gitter, Patrick Godau, Ankit Gupta, et al. Bench-marking and evaluation of ai models in biology: Outcomes and recommendations from the czi virtual cells workshop. arXiv preprint arXiv:2507.10502, 2025.

19. Yair Schiff, Chia-Hsiang Kao, Aaron Gokaslan, Tri Dao, Albert Gu, and Volodymyr Kuleshov. Caduceus: Bi-directional equivariant long-range dna sequence modeling. Proceedings of machine learning research, 235:43632, 2024.

20. Garyk Brixi, Matthew G Durrant, Jerome Ku, Mohsen Naghipourfar, Michael Poli, Gwanggyu Sun, Greg Brockman, Daniel Chang, Alison Fanton, Gabriel A Gonzalez, et al. Genome modelling and design across all domains of life with evo 2. Nature, 652(8112):1349–1361, 2026.

21. Thomas Wolf, Lysandre Debut, Victor Sanh, Julien Chaumond, Clement Delangue, Anthony Moi, Pierric Cistac, Tim Rault, Rémi Louf, Morgan Funtowicz, et al. Transformers: State-of-the-art natural language processing. In Proceedings of the 2020 conference on empirical methods in natural language processing: system demonstrations, pages 38–45, 2020.

22. Peter St John, Dejun Lin, Polina Binder, Malcolm Greaves, Vega Shah, John St John, Adrian Lange, Patrick Hsu, Rajesh Illango, Arvind Ramanathan, et al. Bionemo framework: a modular, high-performance library for ai model development in drug discovery. arXiv preprint arXiv:2411.10548, 2024.

